# scEpiTools: a database to comprehensively interrogate analytic tools for single-cell epigenomic data

**DOI:** 10.1101/2023.04.27.538652

**Authors:** Zijing Gao, Xiaoyang Chen, Zhen Li, Xuejian Cui, Shengquan Chen, Rui Jiang

**Author notes:** These authors are equal contributors (Zijing Gao, Xiaoyang Chen, Zhen Li, Xuejian Cui).

## Abstract

Single-cell sequencing technology has enabled the characterization of cellular heterogeneity at an unprecedented resolution. To analyze single-cell RNA-sequencing data, numerous tools have been proposed for various analytic tasks, which have been systematically summarized and concluded in a comprehensive database called scRNA-tools. Although single-cell epigenomic data can effectively reveal the chromatin regulatory landscape that governs transcription, the analysis of single-cell epigenomic data presents assay-specific challenges, and an abundance of tools with varying types and functionalities have thus been developed. Nevertheless, these tools have not been well summarized, hindering retrieval, selection, and utilization of appropriate tools for specific analyses. To address the issues, we here proposed scEpiTools database with a multi-functional platform (http://health.tsinghua.edu.cn/scepitools). Specifically, based on the comprehensive collection and detailed annotation of 553 articles, scEpiTools groups articles into 14 major categories and 90 subcategories, provides task-specific recommendation for different emphases, and offers intuitive trend analysis via directed graphs, word clouds, and statistical distributions. For single-cell chromatin accessibility data analysis, we proposed a novel ensemble method named scEpiEnsemble, which, along with multiple methods as built-in kernels, can be used for flexible and efficient online analysis via the scEpiTools platform. We envision that scEpiTools will guide tool usage and development for single-cell epigenomic data and provide valuable resources for understanding regulatory mechanisms and cellular identity.

**Author summary:** Compared to single-cell RNA-sequencing data, single-cell epigenomic data can reflect a set of epigenetic modifications at the cellular level. In general, the analysis of these data is typically divided into several steps: 1) retrieving available tools based on the omics of data and tasks; 2) selecting appropriate tools manually; and 3) utilizing the chosen tools to analyze data. However, due to the rapid development of tools and the unique complexity of the data, each of the above steps is extremely challenging for researchers. To provide researchers with great convenience, we developed scEpiTools (http://health.tsinghua.edu.cn/scepitools), a database with multiple functionalities. For instance, given the omics type and the analytic task, researchers can easily browse all the available tools via the hierarchical categorization of scEpiTools, and get recommendation scores from multiple perspectives. Considering that researchers may encounter difficulties in hardware requirements or environment setup, we also provide online analysis with various commonly used tools, as well as a novel ensemble method named scEpiEnsemble. In summary, scEpiTools represents a valuable resource for the single-cell epigenomics community, facilitating retrieval, selection and utilization of appropriate tools for diverse analyses, and helping to drive future advancements in the field.

## Introduction

Recent advances in single-cell sequencing technologies provide significant implications for understanding cellular heterogeneity, developmental biology, and disease mechanisms. To fully exploit the potential of these data, numerous tools have been proposed for upstream and downstream analyses. In single-cell RNA sequencing (scRNA-seq) community, scRNA-tools [1] was proposed to help researchers to navigate the plethora of tools by category. Since its inception, scRNA-tools has been widely used and its updated version further reveals trends in the field with over 1000 collected tools [2], providing a valuable guidance to researchers in selecting tools for analysis.

Unlike scRNA-seq data, single-cell epigenomic data capture the chromatin regulatory landscape that governs transcription. Recent innovations in single-cell epigenomic technologies have enabled the profiling of a wide range of omics data, such as chromatin accessibility, DNA methylation, chromatin interaction, histone modification, and chromosome conformation [3]. The analysis of single-cell epigenomic data has assay-specific challenges such as extreme sparsity and significantly lower sensitivity and higher dimensions, leading to numerous tools with various types and functionalities. Given the constantly evolving landscape of single-cell epigenomic tools and the associated challenges with data complexity and interpretation, it is crucial for researchers to stay up-to-date with these developments to ensure the accuracy and reliability of analyses. Therefore, a comprehensive and intuitive database for interrogating single-cell epigenomic tools is in pressing need. However, establishing a comprehensive database for single-cell epigenomic data analysis poses significant challenges. Firstly, the increasing number of analysis tasks makes scientific categorization of tools complex, necessitating a hierarchical categorization system. Secondly, researchers require task-specific guidance when selecting diverse tools, preferably through an evidence-based recommendation system rather than a simple list of tools. Thirdly, non-methodology papers, such as review studies or those covering sequencing technologies, can provide valuable references and should not be overlooked. Fourthly, the varying configurations and strict environmental requirements of different tools create obstacles to implementation, and the impracticality of local analysis due to the increasing number of profiled cells and complexity of tool usage underscores the need for a more convenient online analysis platform.

To address these challenges, we developed scEpiTools, a user-friendly database with systematic collection and careful annotation of 553 articles (constantly being updated), advanced searching with versatile search filters and sort options, hierarchical browsing with various statistical charts, and custom recommendation for algorithm, review, and sequencing technology. We also conducted trend analyses and statistical analyses for the collected articles. To facilitate the analysis of single-cell chromatin accessibility sequencing (scCAS) data, we proposed a novel ensemble method named scEpiEnsemble and provided it along with multiple built-in kernels on an online analysis platform. In addition, we elaborates extensive application scenarios of scEpiTools for tool selection and benchmarking, and analyzing scCAS data online. Furthermore, scEpiTools provides services such as flexible data downloading and docker images of widely-used tools. We posit that scEpiTools will effectively empower researchers with an all-encompassing comprehension of current research in the field of single-cell epigenomics.

## Design and implementation

### Data collection

We adopted a series of standardized procedures for reliable collection of articles, including methodology articles, reviews, sequencing technologies, and studies. Firstly, a total of 3227 articles with the keywords related to single-cell epigenomics were retrieved from the PubMed, arXiv and bioRxiv (as of Mar 2023). Secondly, the candidate articles were filtered based on their relevance to the field of single-cell epigenomics. Then the references of each article are manually reviewed by at least two independent researchers to check for any missing articles. Ultimately, the number of candidate articles was reduced to 553. The full text and relatively source code of each candidate article was then manually reviewed in detail. The general information such as title, journal, digital object unique identifier (DOI), publication date, citations, reference, publication status, abstract and description were extracted firstly (Fig 1).

**Fig 1.**
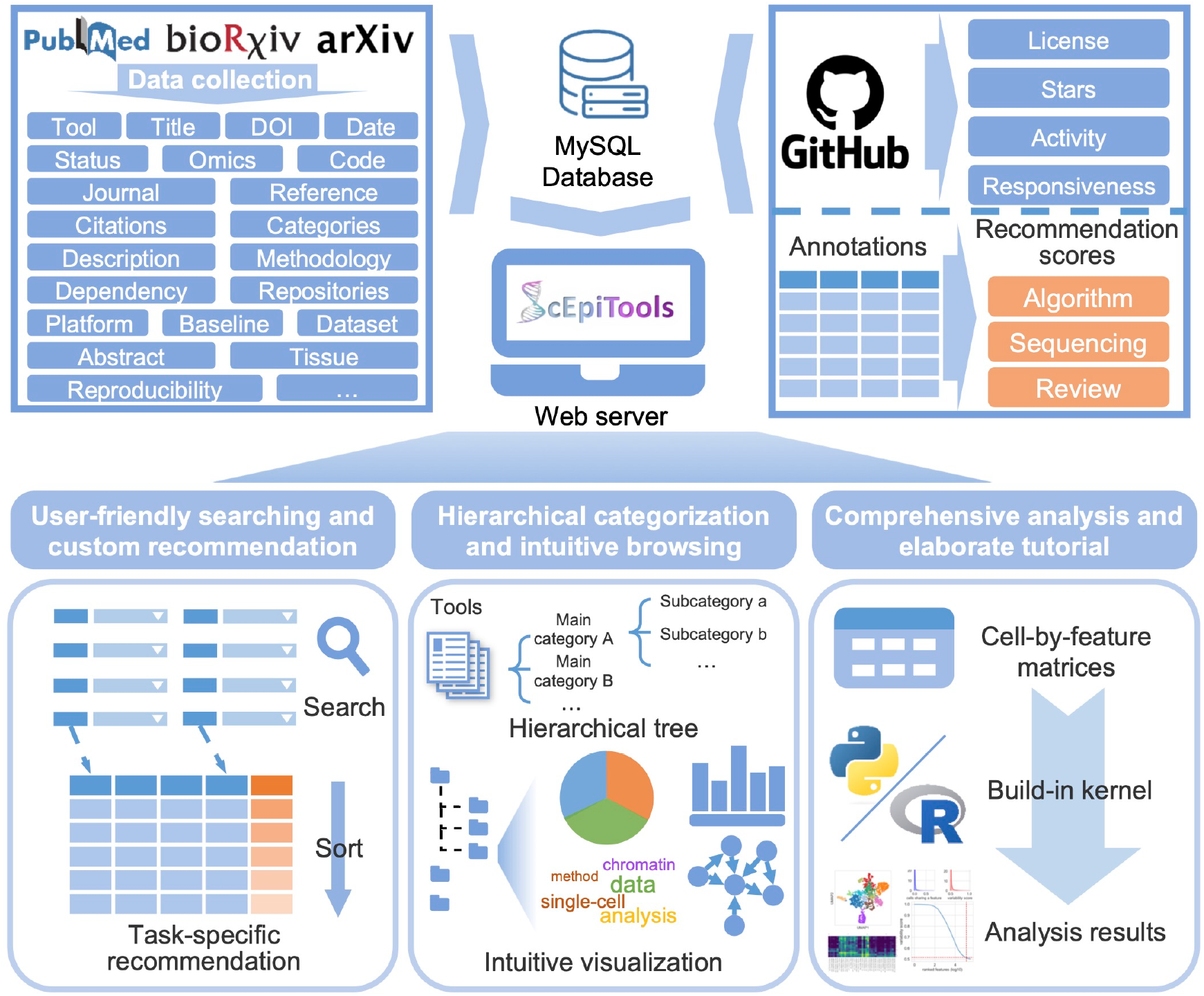
Overview of data collection, processing and annotation, and major features of scEpiTools. scEpiTools is a comprehensive repository comprising 553 epigenomic tools that have been meticulously classified into 14 main categories and 90 subcategories.

### Data processing and annotation

After collecting hundreds of articles, our first step was to categorize them into a hierarchical tree based on their tasks and problems. After summarizing, we categorized them into 14 major categories and 90 subcategories (Table S1). Note that a tool may be applied to multiple different tasks and thus some tools were categorized into multiple categories. For example, SCALE [4] is a deep learning algorithm for dimensionality reduction, but it also provides assistance for imputation of scCAS data by filling missing values and removing potential noise. Therefore, we also categorized it into imputation other than dimensionality reduction.

To facilitate the recommendation and benchmarking of single-cell epigenomic tools, each tool in scEpiTools was annotated manually with extensive details: (1) the GitHub information including stars, activity, responsiveness, and last maintaining date. (2) the dependencies of the software tools, methodologies, baseline methods, number of dataset and figures of the model and source codes. (3) the three novel recommendation scores for three different tasks, encompassing algorithm, sequencing technology and review. These scores are computed via a weighted sum of eight key factors, derived from five distinct aspects, namely influence, usability, publication date, source code maintenance status, and other metrics. (Text S1 and S2).

### Website development and web interface

scEpiTools runs on a Linux-based Apache web server (https://www.apache.org) and utilizes the Bootstrap v3.3.7 framework (https://getbootstrap.com/docs/3.3/) for its web-frontend display. Advanced tables and charts are implemented using plug-ins for the jQuery and JavaScript libraries, including DataTables v1.10.19 (https://datatables.net) and morris.js v0.5.0 (https://morrisjs.github.io/morris.js/index.html), respectively. The backend of the server uses PHP v7.4.5 (http://www.php.net), and all data is stored in a MySQL v8.0.20 (http://www.mysql.com) database. The platform is compatible with the majority of mainstream web browsers, including Google Chrome, Firefox, Microsoft Edge, and Apple Safari.

scEpiTools comprises seven main pages (Text S3). The Home page highlights the key features and major applications of our database. Users can utilize advanced options to find suitable tools and access diverse recommendation scores at the Search page. The Browse page enables users to navigate tools hierarchically for specific tasks and access a range of statistical measures, including abstract word clouds, directed graphs illustrating relationships between tools, and statistical distributions (Text S4). The word cloud presents the primary focus of each category, while the directed graph provides an intuitive representation of comparative relationships among tools within the current category. The Analysis page provides users with the ability to perform scCAS online analysis effortlessly without requiring programming skills with scEpiEnsemble and three other mainstream kernels. At the Download page, users can access detailed annotation information for articles as well as docker images of mainstream methods. The Help page offers instructions on how to use the website, as well as a list of frequently asked questions with corresponding answers. Additionally, we welcome users to contribute any articles or tool methods that may have been overlooked during our collection process on the About page.

## Results

### Overview of the scEpiTools database

The current version of scEpiTools contains 553 single-cell epigenomic tools published or preprint from 2015 to 2023 and is being continuously updated, including 268 articles for tool development, 62 articles for review, and 223 articles for sequencing technologies and applications. Based on the data available in scEpiTools, it was found that as of 2019, the number of tools specifically designed for single-cell epigenomics research was a mere 149. However, as of March 2023, this number has increased nearly four-fold, suggesting the progress made in single-cell sequencing technology and the accumulation of epigenomic data (Fig 2A). Furthermore, on average, these articles also have a considerable citation count, with over 28% of articles having a citation count exceeding 50, as of the last updated date for articles in the database (Fig 3A). The rapid growth in the number of these single-cell epigenomic tools also reflects the increasing research interest among scientists in single-cell epigenomics in recent years.

**Fig 2.**
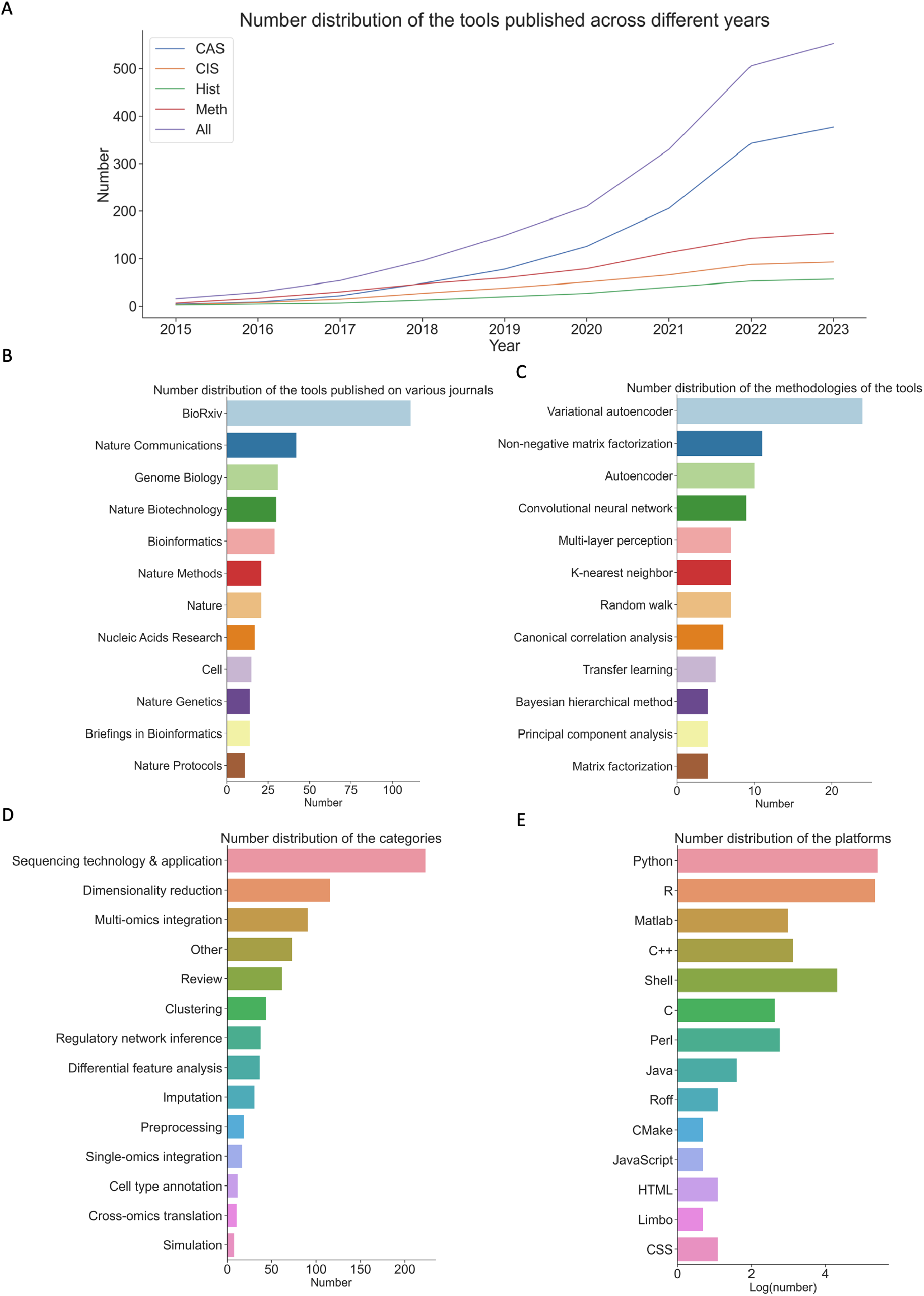
Statistics of single-cell epigenomic tools in scEpiTools. (A) Accumulated number distribution of single-cell epigenomic tools of different omics between 2015 and 2023. The four omics are chromatin accessibility sequence (CAS), chromatin interaction sequence (CIS), DNA methylation (Meth), and histone modification (Hist), and we also considered the total number of articles (All) in each year. (B) Number of tools across 14 main categories. (C) Number of methodologies of the tools (top 12). (D) Number of tools using different platforms. (E) Number of tools published various journals (top 12).

**Fig 3.**
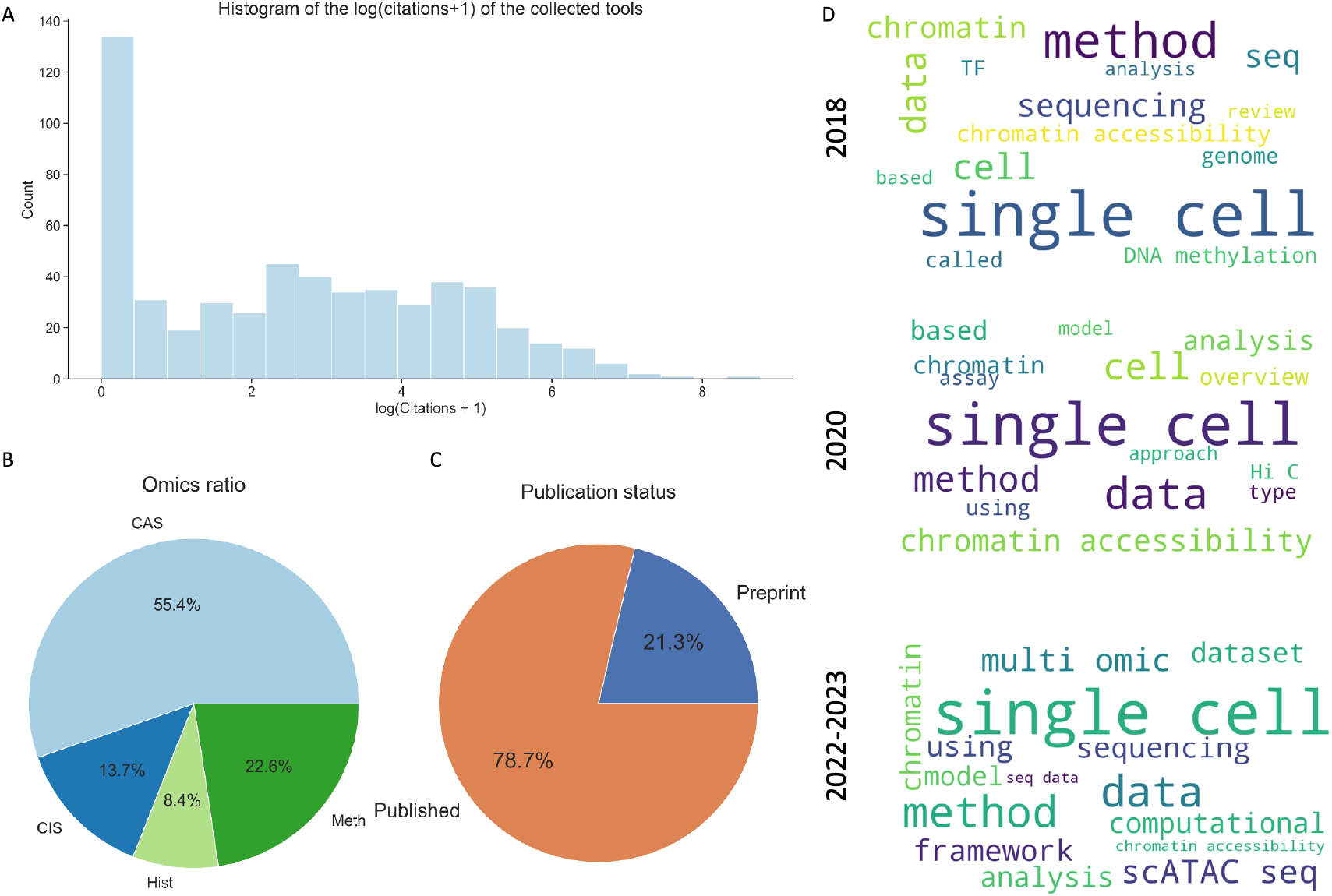
Statistics of single-cell epigenomic tools in scEpiTools. (A) Citation counts for single-cell epigenomic tools with logarithmic transformation. (B) Proportion of different omics in 553 single-cell epigenetic tools. (C) Publication status of 553 single-cell epigenetic tools. (D) Word cloud of the description of the articles published in 2018, 2020 and 2022-2023.

### Statistics and trends of single-cell epigenomic tools

#### Statistics of categories

We have categorized collected tools into 14 main categories based on their types and functionality, including dimensionality reduction, clustering, integration of single-omics data, among others. Throughout the period spanning 2015 to 2023, it has been consistently observed that sequencing technology and application, along with dimensionality reduction tools, have remained the most salient research areas (Fig 2B). This phenomenon can be attributed to the inherently high-dimensional properties of sequencing data such as scATAC-seq. Additionally, it is worth noting that up until 2022, there has been a substantial increase in the number of tools for multi-omics integration, which is indicative of a shift in focus within the field of epigenetics from a single-omics to multi-omics approach, owing to the rapidly development of multi-omics sequencing technologies (Table S2-S3, Fig S1).

#### Statistics of methodologies

We categorized and organized the methodological foundations of methodological articles. Our analysis revealed that the collective utilization rate of variational autoencoder (VAE) [5] and autoencoder (AE) [6] methods surpassed 12.78%, with VAE being the predominant approach (Fig 2C). Noteworthy examples of VAE methods employed in single-cell epigenomics research include scVAEIT [7], SCALEX [8], and GLUE [9]. VAE is a generative model originally developed for image data, but its powerful feature extraction ability can adapt well to single-cell data such as scCAS and single-cell chromatin interaction sequencing (scCIS) data, which inherently contain high noise and high dimensionality. Hence, this might be a potential explanation for the frequent utilization of VAE in single-cell epigenomics data. Furthermore, non-negative matrix factorization [10] and convolutional neural networks are prevalent methodologies for dimensionality reduction or feature extraction in single-cell epigenomic analysis. The prevalence of these approaches underlines the urgent need for addressing the challenge of high dimensionality in single-cell epigenomics analysis.

#### Statistics of platforms

We found that among the collected tools that provide source codes, those developed using Python and R are the most prevalent (Fig 2D), and most methods provide open-source licenses (Fig 4A). This is partly due to the fact that R is one of the most commonly used programming languages in the field of bioinformatics [11] and Python is one of the most widely used language for machine learning [12]. We also observed that with the increasing popularity of deep learning algorithms, there has been a growing trend towards the use of machine learning algorithms in the collected tools, which has led to a higher proportion of Python in single-cell epigenomic analysis tools. In addition, C++, MATLAB and Shell scripting are also popular among these tools. In single-cell epigenomics research, researchers need to frequently process large amounts of sequencing data, which requires high efficiency, convenience, and visualization capabilities for computation. Therefore, these three programming languages are widely used due to their powerful data processing and analysis abilities. Additionally, we also analyzed the GitHub information of the collected articles, based on the observation of GitHub activity, responsiveness, and stars, the tools generally demonstrate satisfactory usability. (Fig 4B-4D, Text S5).

**Fig 4.**
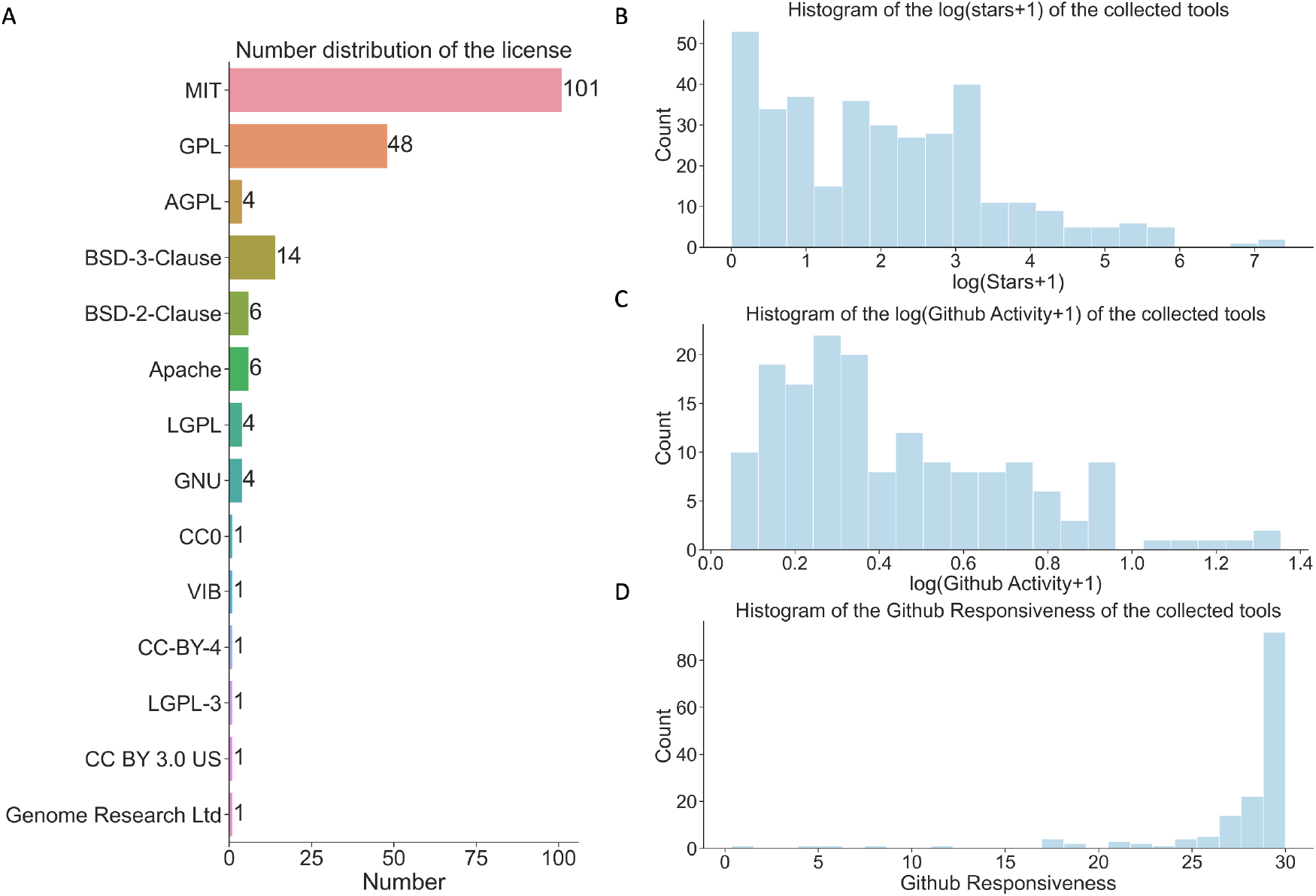
Statistics of single-cell epigenomic tools that open-sourced on GitHub. (A) The distribution of licenses for single-cell epigenetic tools with open-source code on GitHub. (B) GitHub stars with logarithmic transformation. (C) GitHub activity with logarithmic transformation. (D) GitHub responsiveness of the tools.

#### Trends of single-cell epigenomic tools

It is worth noting that among the four types of epigenomic data, methods related to scCAS has consistently been the most abundant each year. In particular, scEpiTools contains 377 articles for chromatin accessibility sequencing, 93 articles for chromatin interaction sequencing, 57 articles for histone modification, and 154 articles for DNA methylation (Fig 3B). This is potentially due to the fact that chromatin accessibility data are closely related to regulatory landscape, cellular development and differentiation, as well as diseases, and also because the ease of obtaining scCAS data compared to other omics data and the availability of numerous sequencing technologies for scCAS data. From 2015 to 2022, there has been a consistent increase in the number of single-cell epigenomic analysis data published each year relative to the previous year. In 2022 alone, 176 tools for analyzing single-cell epigenomic data were published, which represents more than a three-fold increase compared to 54 tools of 2017. This trend underscores the growing importance of such tools for researchers who require them to facilitate investigations of single-cell epigenomic data in the face of rapidly accumulating volumes of data. Additionally, we found that preprints accounted for 21.3% of the total publications, which indicates the rapid development of computational methods and sequencing technologies for analyzing single-cell epigenomic data (Fig 2E, Fig 3C). We selected article descriptions published in 2018, 2020, and 2023, and used them to generate word clouds. We observed that the term “sequencing” was frequently used in 2018, whereas in 2022-2023, high-frequency terms included “deep”, “computational”, and “multi-omics” (Fig 3D). This suggests that with the recent advancement of multi-omics sequencing technologies [3] and the rapid development of deep neural networks, researchers are increasingly focusing on utilizing computational frameworks for the analysis of multi-omics data.

### scEpiTools enables online analysis of scCAS data

scEpiTools provides an intuitive tutorial-style interface, making it accessible to users without coding experience to perform single-cell epigenomic data analysis. Moreover, it assists users in overcoming insufficient local computing resources. The platform offers a complete analysis pipeline, including preprocessing, cell type annotation, and visualization, utilizing built-in kernels of EpiScanpy [13], Signac [14], and SnapATAC [15]. In order to improve the quality of single-cell epigenomic data analysis, we have proposed scEpiEnsemble, an ensemble method that leverages the strengths of EpiScanpy, Signac, and SnapATAC, to obtain comprehensive insights into single-cell epigenomic data analysis. (Text S6, Table S4-S7).

Similar to existing pipelines for analyzing single-cell epigenomic data, scEpiEnsemble initiates by performing a set of optional preprocessing steps such as binarization, quality control, term frequency–inverse document frequency (TF-IDF) transformation in Signac [14], and normalization. As an ensemble method, scEpiEnsemble employs principal component analysis (PCA) from EpiScanpy as well as dimensionality reduction methods from Signac and SnapATAC. We provided three approaches to integrate the results of dimensionality reduction, namely direct concatenation, min-max normalization and z-score normalization. Following the dimensionality reduction, which is essential for downstream analysis of single-cell epigenomic data, we proceeded to cluster and visualize the outcomes. Furthermore, we computed four metrics, namely, adjusted rand index (ARI), adjusted mutual information (AMI), normalized mutual information (NMI), and homogeneity (Homo), to quantify the consistency between the clustering labels and the true cell type labels (Text S6).

We implemented the complete analysis process on human PBMC [16] dataset an example and visualized the results of clustering using uniform manifold approximation and projection (UMAP). scEpiEnsemble achieved an ARI, AMI, NMI, and Homo of 0.483, 0.641, 0.645, and 0.700, respectively, which were at least 3.5%, 1.9%, 1.8%, and 1.3% higher than the baseline methods, demonstrating the effectiveness and superiority of the ensemble strategy (Fig S2). In summary, scEpiEnsemble not only enhances the accuracy of single-cell epigenomic data analysis, but also furnishes an accessible pipeline for users, particularly those without programming expertise.

### Case applications of scEpiTools

scEpiTools has broad applications in areas such as recommendation and tool selection, online analysis, and tool benchmarking. The comprehensive capabilities of scEpiTools enable users to easily navigate and effectively utilize a range of analytical tools and resources, thus improving the efficiency and accuracy of their research. Here, we present two specific application scenarios of scEpiTools, demonstrating how it can assist researchers in selecting tools and analyzing single-cell epigenomic data (Fig S3).

We firstly consider a scenario where algorithm researchers are interested in developing computational tools for the dimensionality reduction of scCIS data. Prior investigation and benchmark of existing state-of-the-art methods are necessary. In this case, they can leverage our database to access state-of-the-art algorithms and obtain initial insights into the underlying principles and strengths and weaknesses of these tools. As an example, by selecting “Chromatin interaction” as the “Omics” option and “Dimensionality reduction” as the “Category” option at the Search page, a list of 19 records for their query can be obtained. Then they can further sort the tools by the recommendation scores for algorithm, GitHub activity, etc. If researchers want to implement methods based on the mainstream programming languages in the field of bioinformatics, i.e. Python and R, they can select these two platforms in the “Platform” option. The results indicate that Galaxy HiCExplorer 3 [17], Higashi [18] and scHiCluster [19] are the most recommended methods. When researchers enter the Details page of Galaxy HiCExplorer 3, they can obtain more information related to the method, such as the required dependencies and the link to source codes. In addition, algorithm developers often need to benchmark their tools against other methods. In such cases, they can easily investigate the existing benchmarks by navigating to the Browse page and selecting the category of “dimensionality reduction”. The network diagram demonstrates that scHiCluster takes various methods as baselines, indicating its novelty and potentially outstanding performance for the current task.

The second scenario involves analyzing scCAS data online. Suppose a user has profiled a set of scCAS data and wants to perform analysis such as dimensionality reduction and differential feature analysis. They can first select one of the four kernels according to the demand, such as diverse input formats and various chromatin regions, and then obtain a detailed notebook that describes the analysis process and results, as well as downloadable results. Taking EpiScanpy kernel as an example, researchers can prepare the input file as the format of AnnData [20] and select the EpiScanpy as the tool for analysis. Then they can view the individual steps of the tutorial before submitting a task. We have provided a set of default parameters that they can modify, such as the clustering method. After submitting a task, the user can obtain a unique task ID, which can be used to view the status and retrieve the results of the task. When the analysis is completed, scEpiTools will provide a detailed notebook and downloadable AnnData-format file. In addition, if the user provides their email address, an email will be sent automatically when the task is completed.

## Availability and future directions

### Availability

The scEpiTools database is publicly accessible through the website at http://health.tsinghua.edu.cn/scepitools, the source code for scEpiEnsemble is freely available on https://github.com/ZjGaothu/scEpiEnsemble and the source code for plotting and analyses is freely available on https://github.com/ZjGaothu/scEpiTools.

### Future directions

Since 2015, the field of single-cell epigenomics has witnessed a remarkable surge in the number of studies, encompassing a range of sequencing technologies, software tools, and related review articles, with an accelerating pace. Our database has diligently collected and meticulously annotated these tools, organized them into distinct categories based on their functionalities and applications, and evaluated their recommendation scores, culminating in the development of an online analysis platform. With the continued accumulation of high-quality sequencing data and the rapid progress of deep learning techniques, we anticipate the emergence of more diverse and advanced tools in the future. To enhance the quality of our database, we will undertake periodic updates and reviews of the listed tools, ensuring their completeness and accuracy, incorporate the most demanded and widely used tools into our online analysis platform, and consider integrating a chatbot system into the new version of scEpiTools, leveraging state-of-the-art language models such as GPT [21, 22], thus facilitating user engagement and improving their experience.

## Supporting information

Text S1. Definition of GitHub activity and responsiveness.

Text S2. Definition of three recommendation scores.

Text S3. Web interfaces of the scEpiTools.

Text S4. Word cloud and the network diagram.

Text S5. Usability of single-cell epigenomics tools.

Text S6. Details of the scEpiEnsemble.

Fig S1. Number distribution of single-cell epigenomic tools of different main categories published between 2015 and 2023.

Fig S2. Clustering performance comparison of scEpiEnsemble and three other mainstream methods.

Fig S3. Case applications of the usage of scEpiTools.

Table S1. The detailed description of each main category.

Table S2. The number of tools under different main categories.

Table S3. The number of tools under different main categories from 2015 to 2023.

Table S4. The user-defined parameters for the turorial of EpiScanpy.

Table S5. The user-defined parameters for the turorial of Signac.

Table S6. The user-defined parameters for the turorial of SnapATAC.

Table S7. The user-defined parameters for the turorial of scEpiEnsemble.

## Acknowledgments

This work was supported by the National Key Research and Development Program of China [2021YFF1200902] and the National Natural Science Foundation of China [62203236 and 62273194].

